## Supplementary figures and images for "Membrane potential modulates ERK activity and cell proliferation"

### Supplemental Figure

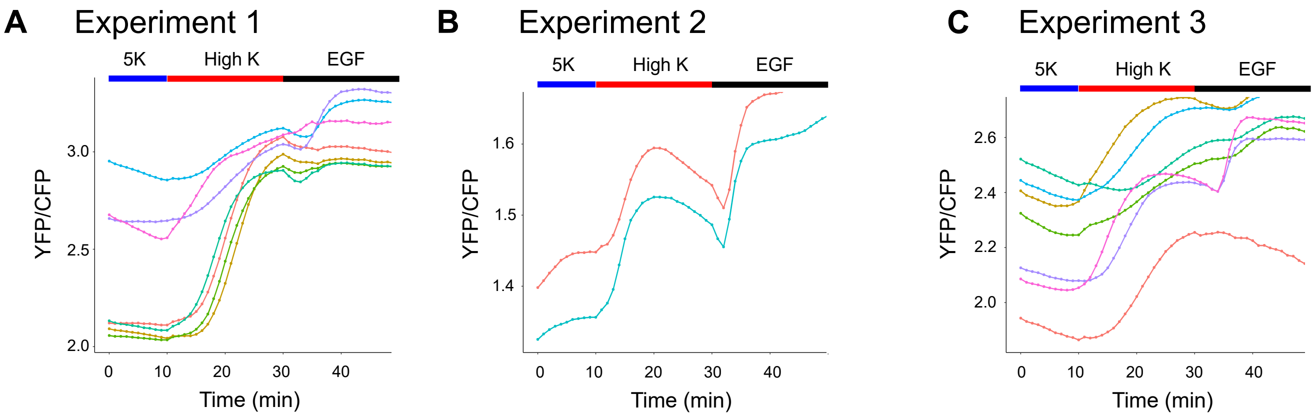

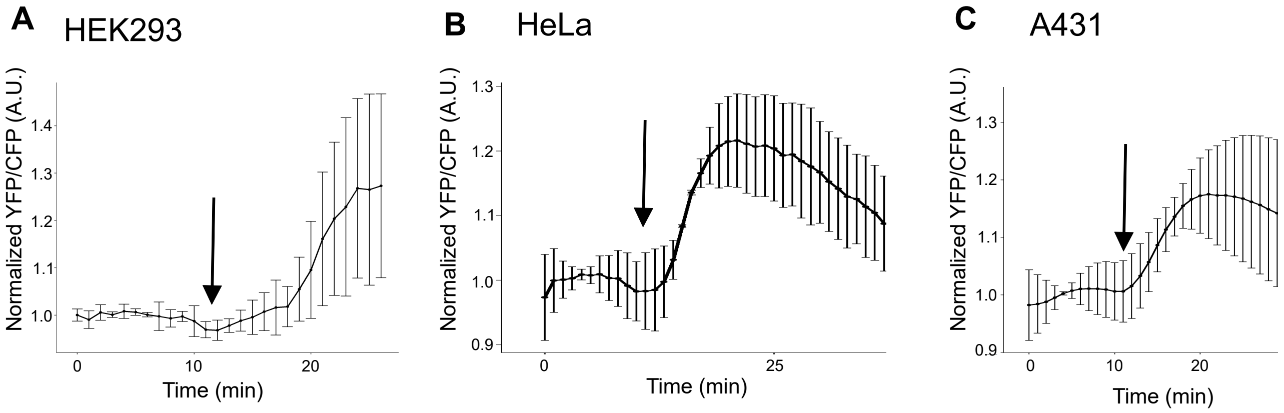

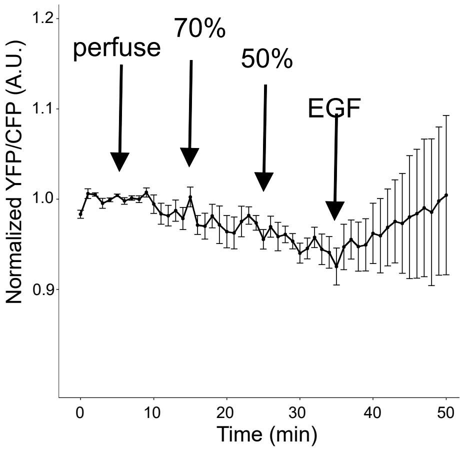
